# Mechanoresponse of epithelial monolayers to in-plane and out-of-plane curvatures imposed by 3D microwells

**DOI:** 10.1101/2022.10.04.510866

**Authors:** Marine Luciano, Marie Versaevel, Eléonore Vercruysse, Sylvain Gabriele

## Abstract

The organization of epithelial tissues with precise spatial definition is essential to various biological processes and to generate curved epithelial structures. However, the regulation of the architecture and dynamics of collective epithelial assemblies by the matrix curvature remains understudied. Here, we photopolymerize microwells of various diameters in hydrogels to form curved epithelial structures such as breast epithelial lobules, and study how in-plane and out-of-plane curvatures modulate the mechanoresponse of epithelial tissues. In-plane curvature governed by the microwell radius drives the centripetal orientation of cells and nuclei close to the edge of the microwell, resulting from contractile forces exerted by a supracellular actomyosin purse-string. Convex out-of-plane curvature imposed at the microwell entrance leads to a vertical orientation of the nuclei towards the microwell axis. We demonstrated that increasing the out-of-plane curvature leads to more flatten and elongated nuclear morphologies with high levels of compacted chromatin. Epithelial cells exhibit higher directionality and speed around the microwell edge, demonstrating that the out-of-plane curvature significantly enhances the cellular trafficking. These findings demonstrate the importance of in-plane and out-of-plane curvatures in epithelial organization and how both can be leveraged to facilitate the engineering of curved structures to study curvature-dependent mechanotransduction pathways.

## 1. Introduction

Cumulative evidence has shown that cells have the ability to sense and respond to changes of the physico-chemical properties of their extracellular matrix (ECM) ^[1]^. Indeed, cells reside in native tissues in a complex microenvironment that guides and regulates their functions ^[2]^. For instance, numerous studies have shown that the mechanical stiffness of the cellular microenvironment has a fundamental effect on cell migration ^[3]^, stem cell differentiation ^[4]^ and can impact tissue regeneration ^[5]^. Recently, evidence is rising that the geometrical properties of the cell’s microenvironment also play an important role in the regulation of cellular functions, such as the intestinal crypt-villus architecture where dividing stem cells reside exclusively in the concave zone ^[6]^.

Previous studies have shown that bidimensional (2D) variations of substrate geometries can affect the spatial organization of stress fibers and focal adhesions, leading to the modulation of internal forces exerted by the actin cytoskeleton on the nucleus ^[7]^. By using protein micropatterns, it has been reported that cells on adhesive islands of constant area, but various geometries exhibited different differentiation profiles. For instance, high aspect ratio micropatterns and high subcellular curvature promoted increased cellular contractility that can lead to osteogenic differentiation of stem cells ^[8]^. It has been shown that cells distinguish between changes of cell-sized positive and negative curvatures in their bidimensional physical environment. Indeed, cells form protrusions at positive curvature zones, whereas they form actin cables at negative ones ^[9]^. It was suggested that these versatile functional structures may help cells sense and navigate their environment by adapting to external geometric and mechanical cues.

In addition to 2D curvature changes, there is increasing evidence that three-dimensional (3D) substrate curvature must be also considered as a relevant material parameter that can influence the cellular organization ^[10]^. Indeed, many epithelial tissues exhibit complex morphologies that are dominated by curved surfaces such as those found in lung alveoli, kidney glomerulus, intestinal villi, and breast acini ^[11]^. Interestingly, it was also reported that tissue curvature and associated changes in apicobasal tension can be considered as fundamental determinants of epithelial tumorigenesis in pancreatic ducts and tubular epithelia of the liver and lung ^[12]^. By using macroscale substrates with radii of curvature from tens to hundreds of micrometers, it was observed that cell alignment and migration in response to 3D cell-scale curvature patterns is a complex process that relies cell-cell interactions and depends on cell types. For instance, isolated vascular smooth muscle cells aligned more weakly than single fibroblasts on cylindrical substrates, whereas both cell types in confluent monolayers aligned prominently with respect to the cylinder axis ^[13]^. It was observed that cells exhibit different morphologies on convex and concave surfaces, to minimize the contact area on concave zones and compressive forces on the nucleus ^[14]^. Furthermore, cell migration speed of human mesenchymal stem cells (hMSCs) was found to be significantly higher on concave spherical surfaces than on convex spherical surfaces, whereas substantial deformations were observed on convex surfaces leading to an increase of lamin A expression in the nuclear envelope ^[15]^.

Although these studies have generated valuable results on the curvature-sensitive cellular response, most of them have been done at the single-cell level and used “rigid” substrates with a non-physiological stiffness ^[13][14][16][17]^. To address this issue, recent works have developed original physico-chemical strategies to pattern hydrogels of tunable rigidities with 3D micropatterns. Interestingly, it has been shown that endothelial cells can self-organize in response to combinatorial effects of stiffness and geometry, independent of protein or chemical patterning ^[18]^. By forming well-defined corrugated hydrogels, it was shown that substrate curvature can affect the thickness of epithelial monolayers and that local curvature controls nuclear morphology and positioning. Together, these works identified the curvature-sensitive cell response as a fundamental capacity of cells but the influence of in-plane and out-of-plane curvatures on multicellular organization in soft anatomically relevant 3D shapes is still unclear.

## 2. Results and Discussion

### 2.1. Microfabrication of bowl-shaped 3D microwells by photolithography in hydro-PAAm hydrogels

Soft hydroxy-polyacrylamide (hydroxy-PAAm) hydrogels ^[19]^ were photopolymerized under UVA illumination from 315 to 400 nm by using the photoinitiator Irgacure 2959 that generates ketyl radicals (Fig. 1a). By illuminating the polymer solution through a transparent chromium optical photomask with dark circular zones, we formed bowl-shaped 3D microwells in soft hydroxy-PAAm hydrogels (Fig. 1b). Hydroxyl groups allowed the hydrogel functionalization with fibronectin (Supplementary Fig. 1), ensuring the establishment of specific cell-substrate adhesions which were required to form a cohesive epithelial monolayer (Fig. 1c) and to avoid cell delamination on curved zones ^[20]^. As shown in Fig. 1d, photopolymerization through an optical photomask allowed to arrange periodically microwells of various sizes over large areas (10×10 mm^2^). We imposed a spacing of 500 μm between two adjacent microwells to minimize interactions between neighbours and to include flat zones as controls. We used circular patterns of 50, 100 and 150 μm in radius that formed microwells of 30 ± 3 μm, 65 ± 3 μm and 100 ± 3 μm in radius (Fig. 1e), which allow to cover the main range of curved geometries found in anatomical structures, such as mammary acini and lobes, intestinal crypts and alveoli ^[21]^. Interestingly, 3D epithelial cavitating spheres of similar sizes were recently reported to spontaneously grow out of an epithelial monolayer during the naive-to-primed transition of human pluripotent stem cells (hPSCs) that model peri-implantation epiblast development ^[22]^. Our technique is robust and leads to highly reproducible microwell morphologies (Supplementary Fig. 2) with a Young’s modulus of 215 ± 20 kPa. The microwell depth was determined by confocal microscopy (Fig. 1f), allowing to establish the relationship between radius and depth of the microwells (Fig. 1g).

**Figure 1.**
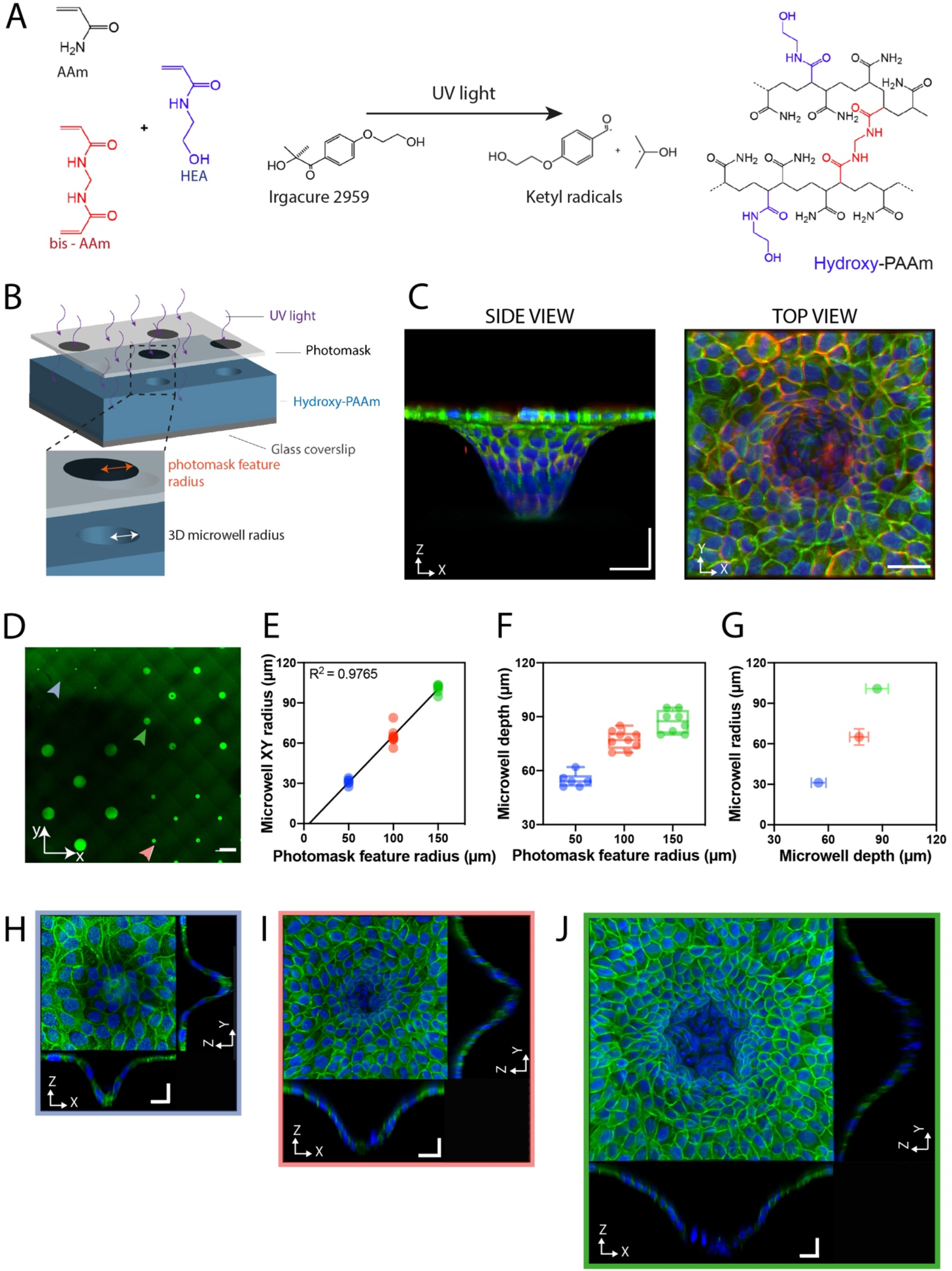
Microfabrication and epithelialization of 3D microwells. (A) The radical polymerization of acrylamide (AAm, in black), bis-acrylamide (bis-AAm, in red) and N-hydroxyethylacrylamide (HEA, in blue), which is amorced by the photoinitiator Irgacure 2959 under UV illumination, led to the polymerization of a polyacrylamide hydrogel with hydroxyl groups (hydroxy-PAAm). (B) The UV photopolymerization of hydroxy-PAAm hydrogels through a chromium optical photomask composed of transparent and circular dark zones formed bowl-shaped 3D microwells in hydroxy-PAAm hydrogel. (C) Confocal images showing side (*xz*) and top (*xy*) views of an epithelial monolayer covering the surface of a 3D microwell. Cadherins are labelled in red, actin filaments in green and DNA in blue. Scale bars are 50 μm. (D) Large image of the spatial distribution of 3D microwells of various diameters in a hydroxy-PAAm hydrogel that was incubated with green fluorescent microbeads. The scale bar is 500 μm. (E) The microwell radius (*xy*) is linearly related to the radius of the photomask patterns (R^2^=0.9765). Evolution of (F) the microwell depth (*xz*) as a function of the photomask feature radius and (G) the microwell radius as a function of the microwell depth. Values related to a pattern radius of 50, 100 and 150 μm on the photomask are in blue, red and green, respectively. Typical confocal images showing top (*xy*) and side (*xz*) and (*yz*) views of 3D microwells of (H) 30 8 3 μm, (I) 65 ± 3 μm and (J) 100 ± 3 μm covered by an epithelial monolayer. Scale bars represent 20 μm.

As shown in Fig. 2, microwells were characterized by a combination of an in-plane (*xy*) curvature, C_x_, determined by the microwell radius, R, as C_x_=1/R, with an out-of-plane (*xz*) curvature C_z_ localized at the microwell entrance and determined as C_z_=1/h. As presented in Fig. 1e, microwells of 30 ± 3 μm, 65 ± 3 μm and 100 ± 3 μm in radius are characterized by an in-plane curvature C_x_ ~0.033 μm^−1^, 0.015 μm^−1^ and 0.01 μm^−1^, respectively. We will first consider the individual effect of the in-plane curvature on epithelial monolayers.

**Figure 2.**
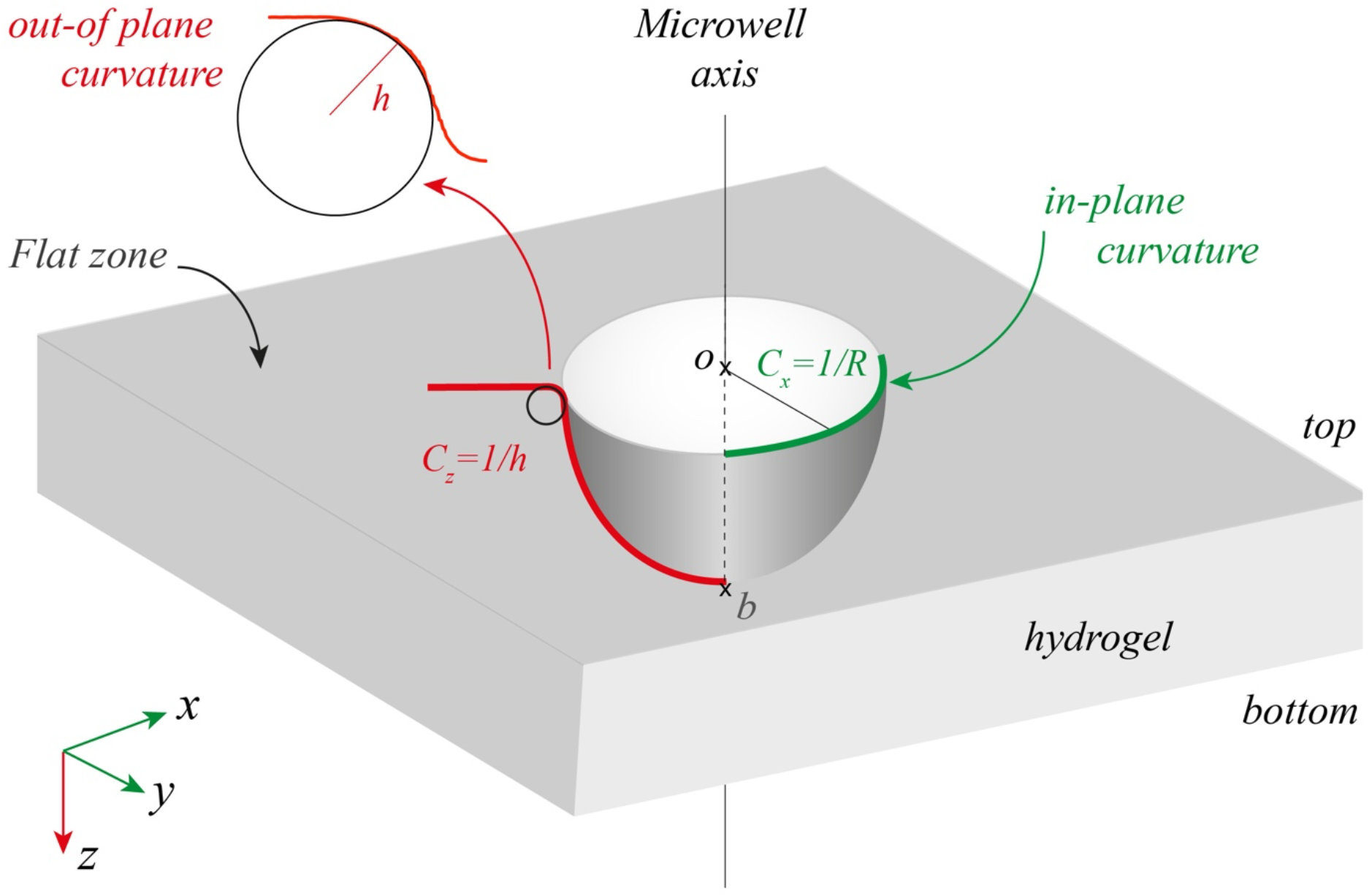
Schematic representation of the in-plane and out-of-plane curvatures in 3D microwells. Microwells formed by photopolymerization in hydrogels are characterized by a combination of an in-plane (*xy*, in green) curvature, C_x_, determined by the microwell radius, R, as C_x_=1/R, with an out-of-plane (*xz*, in red) curvature C_z_ localized at the microwell entrance and determined as C_z_=1/h. A flat hydrogel zone surrounds each 3D microwell.

### 2.2. Cytoskeletal and nuclear orientation occurs around the edge of a 3D microwell and depends on its in-plane curvature

Our results confirmed that Madin-Darby canine kidney (MDCK) epithelial cells organized into a uniform monolayer in and around microwells of 30 μm (Fig. 1h, Supplementary Movie S1), 65 μm (Fig. 1i, Supplementary Movie S2) and 100 μm (Fig. 1j, Supplementary Movie S3) in radius after 24 hours in culture without any delamination. A closer look of the architecture of the epithelial monolayer around the edge of the microwells indicated significant changes of the cell orientation. To get further insight into these changes we used a MDCK cell line expressing red fluorescent E-cadherin at cell-cell contacts. As shown in Fig. 3a, epithelial cells around the microwell edge orientated their long axis towards the (*xy*) center of the microwell. Our results indicated that the cell angle between the long cell axis and the line passing through the cell center of mass and the microwell center was larger for high in-plane curvature (36.6± 20.1° for 0.033 μm^−1^) than for low in-plane curvature values (12.7± 8.9° for 0.01 μm^−1^, Fig. 3b), demonstrating that low in-plane curvature imposed by larger microwell diameters leads to a higher centripetal orientation of the epithelial cells localized around the microwell edge.

**Figure 3.**
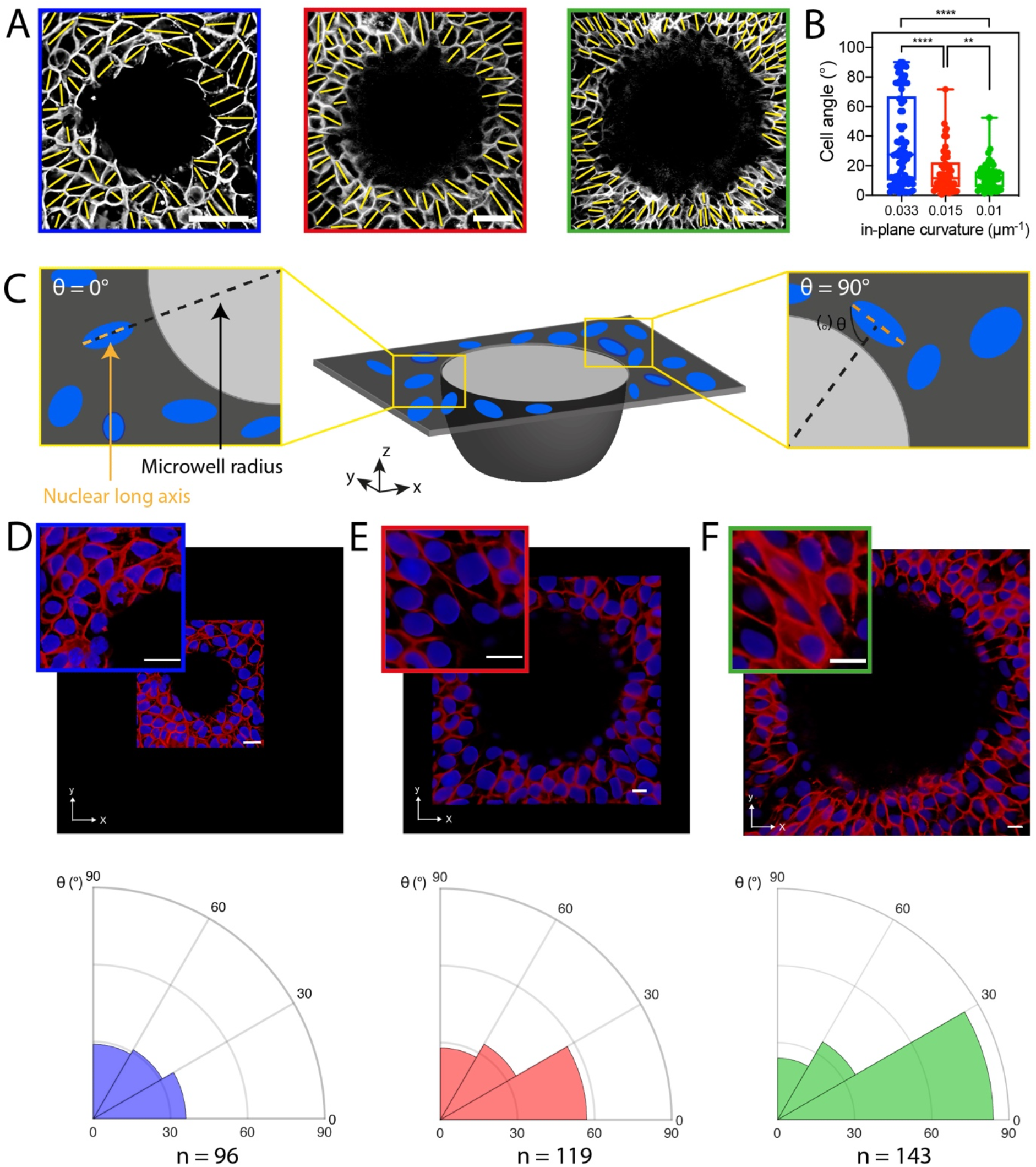
In-plane curvature modulates cell and nuclear orientation at the microwell edge. (A) Cadherin immunostained images were used to determine (B) cell orientations at the edge of 30 μm in radius (n=89, in blue), 65 μm in radius (n=68, in red) and 100 μm in radius (n=75, in blue) microwells (n=3 for each condition). The orientation of the long cell axis is depicted by a yellow segment. Scale bars are 50 μm. (C) Schematic representation of the nuclear orientation within the flat epithelial tissue located around the edge of 3D microwells. The nuclear orientation was characterized by the angle θ formed between the microwell radius and the nuclear long axis. Angles ranged from θ = 0° for nuclei parallel to the radius (left box) to 8 = 90° for nuclei oriented perpendicular to the radius (right box). Confocal images in the (*xy*) plane and corresponding radial orientation of nuclei around the edge of microwells of (D) 30 μm in radius (n=96, in blue), (E) 65 μm in radius (n=119, in red) and (F) 100 μm (n=143, in green) in radius. Cadherins are labeled in red and nuclei in blue. Scale bars are 20 μm. **p < 0.002 and ****p < 0.0001.

Based on the effect of the microwell diameter on the cell orientation, we next sought to examine the nuclear orientation. Nuclei stained with Hoechst 33342 and localized within a circular region of interest of 60 μm wide around the edge of the microwells were thresholded and an ellipse was adjusted to the nuclear contour to determine the angle between the long nuclear axis and the *xy* microwell axis (Fig. 3c). Our results showed that nuclei around the edge of microwells of 30 μm in radius (Fig. 3d) exhibited a random nuclear orientation, whereas increasing the microwell radius to 65 μm (Fig. 3e) and 100 μm (Fig. 3f) induced a preferential orientation of the nuclei towards the (*xy*) microwell axis, in agreement with the centripetal cellular orientation observed for low in-plane curvatures.

### 2.3. The nuclear orientation around the edge of the microwells relies on contractile actomyosin forces exerted by an actin ring

Knowing that orientation and deformation of the nucleus were related to the amount of contractile tension in actin filaments ^[7][23]^, which is transmitted to the nuclear envelope through LINC complexes ^[24]^, we thus hypothesized that nuclear orientation at the microwell entrance could be due to changes in organization and contractility of the actomyosin network driven by the microwell radius. Confocal imaging performed within a focus plane close to the edge of the microwells (Fig. 4a) clearly indicated the presence of a well-defined supracellular actin structure in 65 μm (Fig. 4b) and 100 μm (Fig. 4c) microwells, that extended as a ring along the edge perimeter. Interestingly, no well-defined actin ring was observed in microwells of 30 μm in radius. Previous studies reported actin ring structures in various developmental and physiological processes ^[25]^ and have established that contractile actin cables connected to a myosin structure can form at high concave edge of lamellipodia, strengthening the hypothesis that specific contractile actomyosin structures can form in response to the curvature imposed at the microwell entrance.

**Figure 4.**
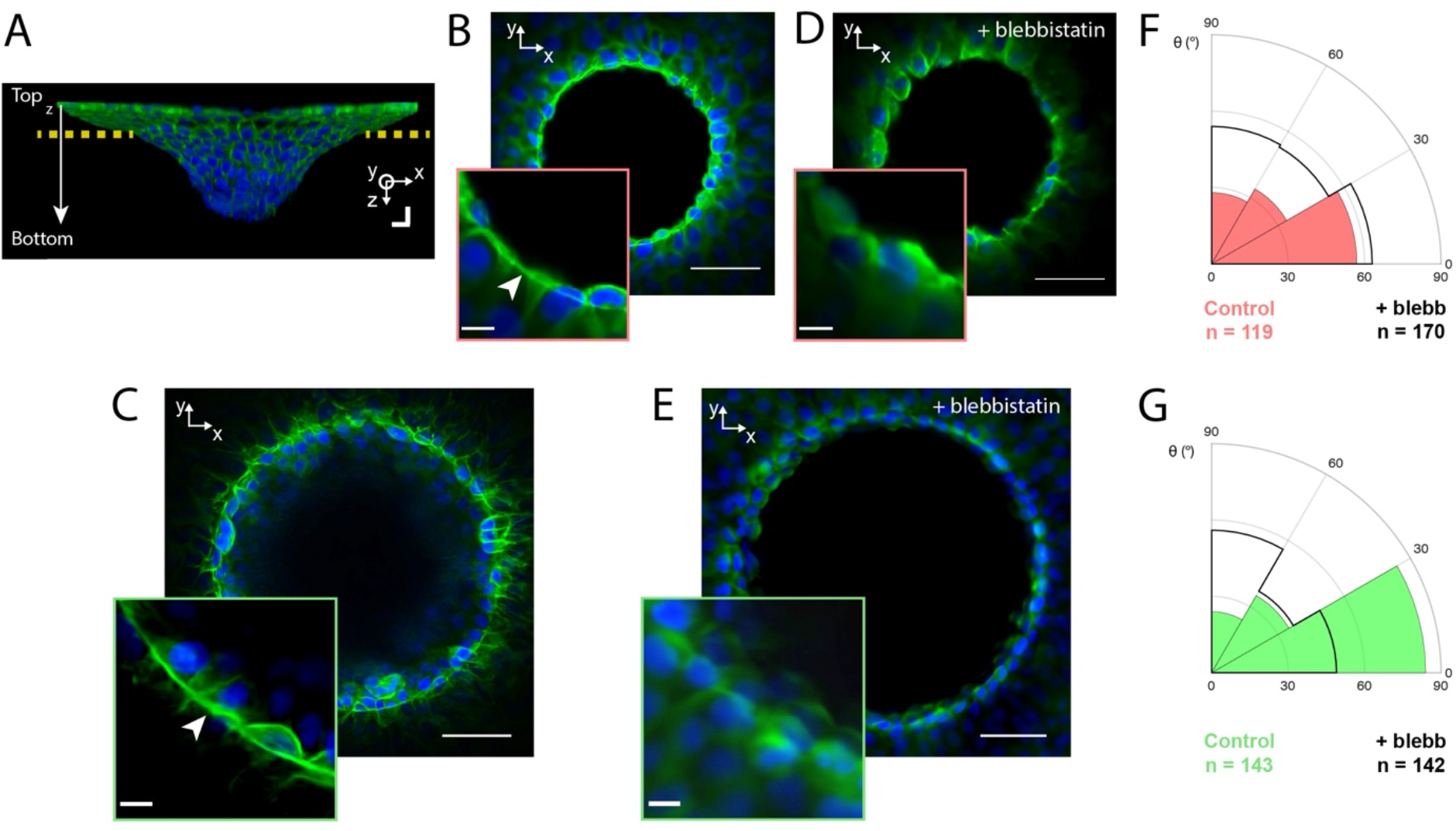
The nuclear orientation around the edge of the microwells relies on contractile actomyosin forces exerted by an actin ring. (A) Confocal side-view (*xz*) of a 3D microwell covered with an epithelial monolayer stained for F-actin (in green) and DAPI (in blue). The yellow dotted line represents the Z plane corresponding to the zone just below the edge of the microwell. Confocal top view (*xy*) showing the presence of an actin ring in microwells of (B) 65 μm and (C) 100 μm in radius, whereas no actin supracellular structure was observed after blebbistatin treatment on microwells of (D) 65 μm and (E) 100 μm in radius. Nuclear orientation for normal (plain colors) and in blebbistatin-treated cells (superimposed in black) localized at the edge of microwells of (F) 65 μm and (G) 100 μm in radius. Values for microwells of 65 μm (n=170) and 100 μm (n=142) in radius are in red and green, respectively.

We tested this hypothesis by altering the myosin II-dependent contractility of the actin cytoskeleton with blebbistatin that blocks the myosin heads and inhibits myosin ATPase activity, suppressing actomyosin contraction ^[26]^. We found that blebbistatin treatments disorganized the actin cytoskeleton on 65 μm (Fig. 4d) and 100 μm (Fig. 4e) microwells and led to the collapse of the actin cable in both microwell sizes. Our findings showed that the disruption of the actomyosin contractility prevented the centripetal nuclear orientation, leading to random nuclear orientations in 65 μm (Fig. 4f) and 100 μm (Fig. 4g) microwells, as observed in non-treated 30 μm microwells (Fig 3d).

Altogether, our findings demonstrate that the fraction of oriented nuclei increases as the in-plane microwell curvature decreases and support the hypothesis that decreasing the in-plane microwell curvature promotes the formation of a contractile actomyosin ring that exerts centripetal forces on cells localized around the microwell edge.

### 2.4. Cells at the edge of a microwell exhibit large vertical nuclear elongation

Based on the modulation of the cellular and nuclear orientation in response to the in-plane curvature, we next sought to determine whether the nuclear organization could be affected by the convex out-of-plane curvature C_z_ (See Fig. 2). To address this question, we first used confocal microscopy at high magnification to obtain the (*xz*) profile (Fig. 5a) of the 30 μm, 65 μm and 100 μm in radius microwells, which were binarized and skeletonized (Fig. 5b) to determine the curvature at each profile point and then the maximal curvature zone (Fig. 5c).

**Figure 5.**
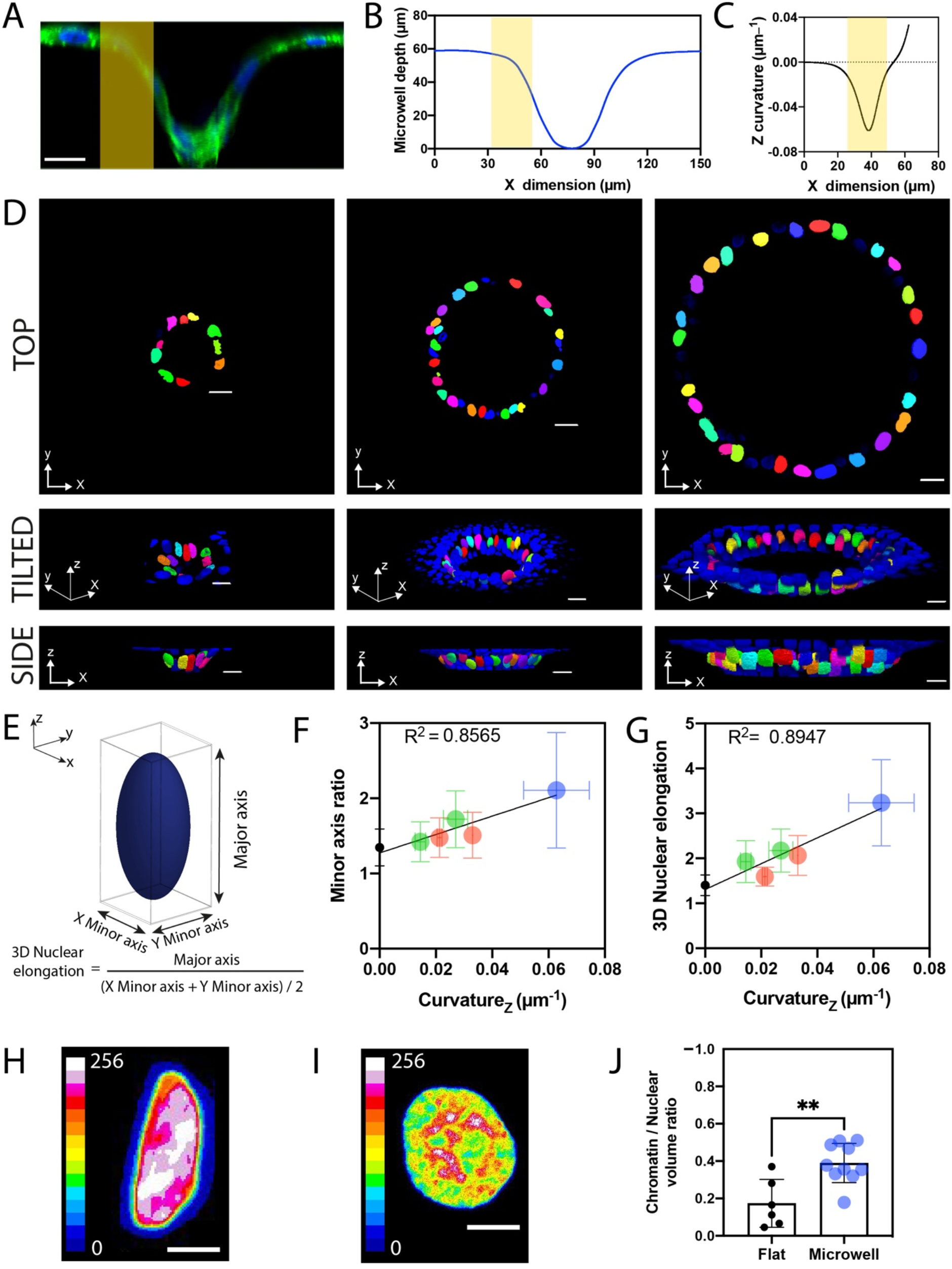
The convex curvature imposed at the edge of a microwell induces a vertical nuclear elongation towards the microwell bottom. (A) Confocal side-view (*xz*) of a 3D microwell covered with an epithelial monolayer. Actin filaments are labelled in green and DNA in blue. Scale bar is 20 μm. (B) Typical outline of a 3D microwell profile obtained after binarization and skeletonization of the confocal side-view. (C) Out-of-plane (*xz*) curvature calculated at each point of the microwell profile presented in (B). The yellow domain in (A-C) corresponds to the maximal convex curvature zone. (D) Top (*xy*), tilted (*xyz*) and side (*xz*) confocal views of nuclei located at the maximal convex curvature zone of microwells of 30 μm, 65 μm and 100 μm in radius. Nuclei were thresholded and color-coded for better spatial representation. Scale bars are 20 μm. (E) Schematic representation of an elongated nucleus with the major axis oriented towards the microwell axis. (F) The minor axis ratio and (G) the 3D nuclear elongation were both linearly related with the maximal vertical curvature of microwells of 30 μm (in blue), 65 μm (in red) and 100 μm (in green) in radius. Black points correspond to the nuclear elongation observed on flat zones. Typical images of the chromatin condensation obtained from a Z-projection for nuclei localized in (H) the maximal curvature zone and (I) a flat zone of a 30 μm microwell. Intensities of DNA staining were digitized in 256 bits and color coded for each Z-stack. Highly condensed domains show higher fluorescence intensity (white zones) with respect to the less condensed ones (blue zones). (J) Chromatin to nuclear volume ratio for nuclei localized on a flat zone and on a convex curvature zone. **p < 0.0025.

We focused our attention to the focal planes corresponding to this maximal curvature zone for determining the 3D morphology of the nuclei of epithelial cells that interacted with the highest convex curvature zone. As observed on top, tilted and side views, nuclei were homogeneously distributed along the microwell periphery, and their long axis were mainly oriented vertically towards the microwell bottom (Fig. 5d). While we found that nuclei around the microwell edge were characterized by a centripetal organization depending on the in-plane curvature value, this observation clearly indicated that nuclei at the maximal out-of-plane curvature zone were mostly oriented vertically towards the microwell axis.

We then investigated whether nuclei underwent morphological changes in response to the modulation of the out-of-plane curvature. By assuming the nuclear morphology as a 3D ellipsoid with its long axis oriented towards the microwell bottom, we quantified the two other minor axes of the ellipsoid from thresholded nuclei (Fig. 5e). Our findings showed that the minor axis ratio increased linearly with the maximal out-of-plane curvature (Fig. 5f), indicating that increasing the convex out-of-plane curvature led to more flattened nuclei, in agreement with previous observations on the crests of corrugated hydrogels ^[20]^. In addition to be flattened, nuclei significantly elongated as the out-of-plane curvature increased (Fig. 5g). Indeed, our findings showed that the 3D nuclear elongation was linearly related to the out-of-plane curvature C_z_, with a nuclear elongation of ~3 observed on the largest out-of-plane curvature value of ~0.06 μm^−1^. Taken together, our findings demonstrated that large out-of-plane curvatures imposed a vertical orientation of the nuclei towards the microwell axis and induced significant 3D nuclear elongations in epithelial cells.

### 2.5. Elongated nuclei at the maximal out-of-plane curvature zone show high level of chromatin condensation

Considering that nuclear shape changes could be associated with a modulation of chromatin organization ^[7][23]^, we next investigated whether nuclear elongations observed at the convex zone of the microwell entrance could be associated with differences of chromatin state. To gain further insight into the role of the curvature, we focused our attention on the more elongated nuclei corresponding to the largest out-of-plane curvature of ~0.06 μm^−1^ (Fig. 5g). Using a quantitative procedure based on DAPI staining and 3D confocal imaging at high-resolution, we determined the average spatial density corresponding to the ratio between the integrated fluorescence intensity and the volume of the nucleus, which is a reliable indicator of the average chromatin condensation. Our results showed that elongated nuclei localized at the maximal out-of-plane curvature zone exhibited higher levels of fluorescence intensity (Fig. 5h) than rounded nuclei in flat zones (Fig. 5i). Altogether, our findings indicate that elongated nuclei at the out-of-plane curvature zone were characterized by higher chromatin compaction values (0.39 ± 0.11, Fig. 5j) than nuclei in flat zones (0.17 ± 0.13), suggesting that the out-of-plane curvature imposed at the microwell entrance leads to elongated nuclei with high levels of condensed chromatin.

### 2.6. Collective migration is enhanced at the maximal curvature zone of 3D microwells

The morphological and functional consequences of in-plane and out-of-plane curvature changes raise important questions about the dynamics of epithelial monolayers grown on microwell-shaped substrates. Indeed, it is not clear whether cells located at convex curvature zones are replaced by neighbours in response to a dynamic remodeling of the epithelial tissue or hold in place. Recent evidence has shown that epithelial monolayers can adapt to curvature changes to maintain homeostasis ^[20][27]^ but the specific contribution of biologically relevant topographical features such as cell-scale curvature on collective dynamics is still unclear. To address this question, we studied by time-lapse microscopy the displacement of epithelial cells in confluent monolayers of similar densities located on flat hydrogel zones (Fig. 6a) and on a convex zone (Fig. 6b, Supplementary Movie S4) of C_z_~0.025 μm^−1^. Cell tracking experiments were limited to the maximal convex curvature zone (see Fig. 4a-c), partly due to the microwell dimensions that prevent optical observations in the bottom zone of the microwells. Nuclei were labeled with Hoechst 33342 and nuclear displacements were tracked over time to determine the straightness (Fig. 6c) and the speed of each epithelial cell. As shown in Fig. 6d, our findings showed that the straightness of epithelial cells was significantly higher at the convex zone (0.57±0.20) compared to flat zones (0.39±0.18), demonstrating that the out-of-plane curvature imposed at the microwell entrance led to a significant increase of the cell directionality. Interestingly, our results indicated that the cell speed was statistically faster on the convex zone than on flat zones (Fig. 6e), suggesting the convex curvatures can enhance the cellular trafficking.

**Figure 6.**
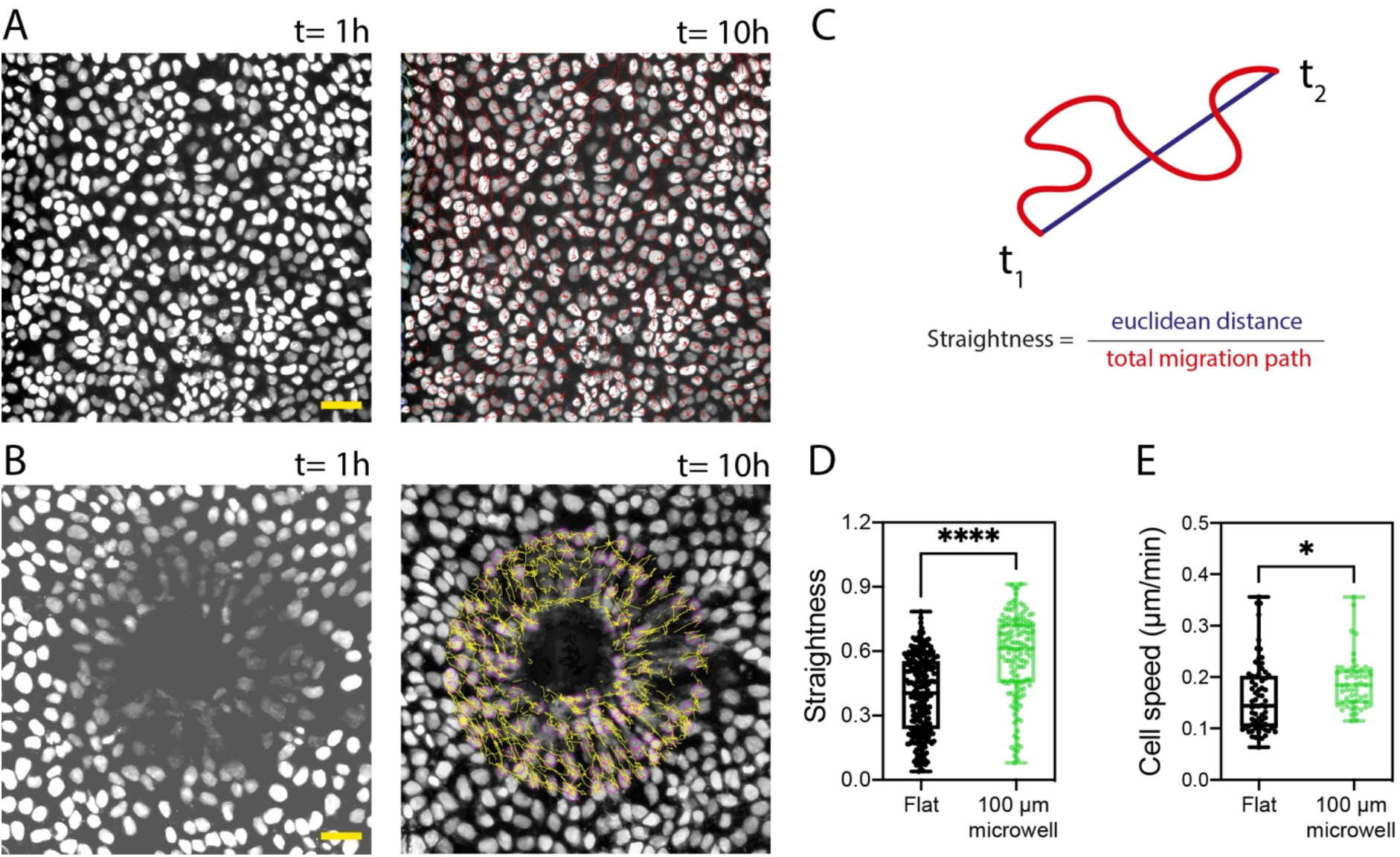
Convex curvature at the microwell entrance enhances collective migration straightness and speed. Time-lapse sequence in epifluorescent mode of the cumulative trajectories over 10 hours of individual cells within a confluent epithelial tissue grown on (A) a flat zone and (B) around the edge of a 3D microwell of 100 μm in radius. Scale bars represent 50 μm. (C) Schematic representation of the straightness parameter, defined by the ratio between the Euclidean distance and the total migration path. (D) Straightness and (E) cellular speed within a confluent epithelial tissue on a flat zone (in black, n=253) and around the edge of a 3D microwell of 100 μm in radius (in green, n=137) with *p < 0.05 and ****p < 0.0001.

Recent advances in hydrogel synthesis and microscale patterning have led to new advances in understanding the role of physical cues of the matrix in cell organization ^[18]^. Among the role of stiffness and small-scale topographies, cumulative evidence has shown that matrix curvature affects spatiotemporal organization of cells and tissues ^[10]^. However, despite rapid advances in microfabrication techniques, including micro-machining ^[17]^, soft lithography ^[28]^, and two-photon polymerization ^[29]^, the robust fabrication of curved cellular environments in soft matrices still remains a difficult task. New technologies are thus required to facilitate more in-depth studies of cell response to the various types of substrate curvatures.

Here we present a robust method to produce in a controlled and reproducible way soft 3D microwells of different sizes and geometries that mimic physiological cavities such as the breast lobules ^[30]^ and kidney Bowman’s capsules ^[31]^. Using these 3D microwells, we report that in-plane (*xy*) and out-of-plane (*xz*) curvatures imposed at the entrance of hydrogel microwells modulate in different ways architecture, nuclear deformations and dynamics in epithelial tissues. Indeed, we showed that in-plane curvatures controlled by the microwell diameter at cellular length scales can drastically influence the self-organization of epithelial cells localized close to the edge of the microwell. Indeed, lowering the in-plane curvature – i.e. increasing the microwell diameter – promotes a centripetal orientation of cells and nuclei towards the (*xy*) microwell center. Furthermore, we demonstrated that lowering the in-plane curvature promotes the formation of a supracellular actomyosin purse-string that exerts contractile forces involved in cellular and nuclear orientation. Interestingly, we found that nuclei of epithelial cells located at the maximal out-of-plane curvature zone were flattened and elongated vertically towards the microwell axis. We demonstrated that the nuclear deformations imposed by the out-of-plane curvature resulted in high level of chromatin compaction suggesting important consequences on the genome integrity. Finally, we showed that out-of-plane curvature not only affected nuclear shape and chromatin organization but also significantly enhanced the cellular dynamic around the convex curvature zone at the microwell entrance.

## 3. Conclusions

In this study, we investigated the influence of in-plane and out-of-plane curvatures changes on the cellular and nuclear morphologies and the subsequent effect on chromatin compaction and migration of epithelial cells in confluent monolayers. Our observations indicate that in-plane and out-of-plane curvatures imposed by the microwell geometry can impact the organization and dynamics of the epithelial tissue and modulate the cytoskeletal forces acting in the nucleus, leading to significant changes of nuclear orientation and function. Based on these new insights of how curvature at the microwell entrance influences epithelial tissues, our findings might contribute to a better understanding of the curvature-responsive mechanical regulation of epithelial cells and provide a robust approach to guide the patterning and self-organization of epithelial cells on curved substrates. Our approach opens a new avenue to create complex microenvironments to investigate fundamental questions in cellular biology such as the influence of the matrix curvature in the regulation of stem cell differentiation and the etiology of diseases associated with curved microenvironments, such as cancer progression ^[12]^. Based on photopolymerization, our strategy allows reproducible and cost-effective hydrogel patterning, can be easily implemented in any laboratory to create customizable shapes of microwells in soft culture matrices and does not require replica molding nor advanced skills in material sciences. Altogether, our findings demonstrate the importance of in-plane and out-of-plane curvature changing in complex cellular organization and show that both can be leveraged to facilitate the engineering of curved structures and study curvature-dependent mechanotransduction pathways. We anticipate that the incorporation of spatially controlled surface curvature as a material design parameter could provide an easy but potentially powerful tool for improving tissue regeneration, biomaterial design, culture conditions and drug testing.

## 4. Experimental Section

### Photopolymerization of the microwells

Round glass coverslips of 22 mm in diameter were incubated 5 min in NaOH 0.1M, then rinsed 3 times with distilled water and dried with a nitrogen flow. A solution of 3-(trimethoxysilyl) propyl acrylate was incubated on clean coverslips for 1 hour. Coverslips were rinsed abundantly with distilled water and dried. A mix solution was prepared containing hydroxyethylacrylamide (HEA), 28.6 % w/w aqueous acrylamide solution, 1.96 % w/w aqueous bisacrylamide solution, 1-[4-(2-hydroxyethoxy) phenyl] - 2-hydroxy-2-methyl-1-propanone (Irgacure 2959, 5mg/ml) and water in order to get final concentrations of 15% and 1/23 ratio of bisacrylamide to acrylamide. The solution was degassed during 15 min by nitrogen bubbling. A 50 μl droplet of the mix hydrogel solution was deposited on each pattern of a quartz photomask (Compugraphics, UK) and squeezed with a circular glass coverslip. The photomask was then insulated with UVA lights (Dymax) at a power of 10 mW/cm^2^ for 10 minutes. Each glass coverslip covered with a microstructured hydrogel was then gently peeled off the photomask and swelled in sterile water for 24 hours. Hydrogels were sterilized by a germicide UV treatment of 15 min and incubated with 75 μg/ml solution of fibronectin for 1 hour and then rehydrated before cell seeding in sterile medium at 37°C during 4 hours.

### Cell culture

RFP E-cadherin expressing epithelial cells from the Madin–Darby canine kidney (MDCK) cell line were cultured in DMEM (Dubelcco’s Modified Eagle’s medium) supplemented with 10% FBS (Fetal Bovine Serum, AE Scientific) and 1% antibiotic (Penicillin, Streptomycin, AE Scientific). MDCK cells were incubated at 37°C with 5% of CO_2_ and in an environment saturated in humidity and seeded on the polyacrylamide hydrogels at a concentration of 100,000 cells/cm^2^.

### Immunostainings and drug treatment

Cells were fixed with a 4% solution of paraformaldehyde and 0.05% Triton X-100 in PBS for 15 min at 37°C and washed three times in PBS. Nuclei and actin cytoskeleton were labelled by an incubation for 1 hour at RT with Hoechst 33342 and Alexa Fluor 488 Phalloidin (Molecular Probes, Invitrogen) and then rinsed 3X with PBS. The coverslips with hydrogel niches were then placed on a microscope sample-holder and covered with glycerol for imaging. Actomyosin contractility of MDCK cells was inhibited with 100 μM of blebbistatin for 1 hour. Cells were then immediately fixed and stained.

### Confocal imaging

Time-lapse and fluorescent images of immunostained tissues in microwells were taken with an inverted confocal microscope (Nikon TI2 A1R HD25, Japan) with a high-resolution mode, a large field of view (25 mm), 3 axes of motorization, 4 lasers and LEDs. High resolution optical silicon lenses (x25, x40) were used for Z acquisitions. Time-lapse acquisitions were performed with 10 minutes time intervals for a total duration of 10 hours by using a cage incubator (Okolab) to maintain the temperature at 37°C, 5% CO_2_ level and a high level of humidity.

### Microwell profiles

Samples were incubated with a solution of fluorescent particles of 0.2 μm in diameter (Fluospheres, Molecular Probes, Eugene, OR) diluted in glycerol. Confocal stacks were performed to obtain the transversal (*xz*) cross-section. Microwell profiles were obtained from the outline of the niches topography thanks to binarization and skeletonizing of transversal (*xz*) cross-sections. A hyperbolic tangent function, y = tanh (x) was then fitted to the half of the microwell profile by a non-linear regression. The curvature was calculated at each point on the fitted equation by κ = ÿ / (1+ ý2) 3/2 where the dot and double dot represent the first and the second derivative of the function, respectively. The local maxima and minima of curvature correspond to the most convex and most concave points, respectively. The quantification was performed from at least 3 cross-sections obtained from 3 independent replicates.

### 2D and 3D nuclear orientation

A Maximal Intensity Projection (MIP) image was created from the Z slices corresponding to the top of the microwells. A circular region of interest (ROI) centered on the center of the niche and having a radius of 60 microns larger than the niche radius was selected on each MIP image. Nuclei entirely included in the ROI were fitted by an ellipse defined by a long and a short axis. The angle formed between the long nuclear axis and the axis passing through the microwell center and the ellipse center of mass was used to determine the nuclear orientation. 3D nuclear orientations and elongations were obtained from a 3D thresholding of Z stacks in NIS Elements. Nuclei included in the flat region around the niches (for control) or in the 30 first microns in depth at the entry of the niche were selected at their 3D pitch as well as 3D elongation were measured.

### Mechanical measurements

The Young’s modulus of immersed microwells was measured with indentation (Chiaro, Optics11, The Netherlands) using spherical probe with a radius of 9 μm and a stiffness of 0.49 N/m. Young’
ss moduli were determined by fitting force-indentation curves with the Hertz equation.

### Statistical analysis

Differences in means between groups were evaluated by two-tailed Student’s *t*-tests performed in Prism 9 (GraphPad Software). For multiple comparisons, the differences were determined by using an analysis of variance followed by Tukey’s post-hoc test. *p < 0.0500; **p < 0.0100; ***p < 0.0010; ****p < 0.0001; n.s., not significant. Unless stated otherwise, all data are presented as mean ± standard deviation (s.d.).

## Supporting Information

Supporting Information is available from the Wiley Online Library or from the author.

## Acknowledgments

S.G. acknowledges funding from FEDER Prostem Research Project no. 1510614 (Wallonia DG06), the F.R.S.-FNRS Epiforce Project no. T.0092.21 and the Interreg MAT(T)ISSE project, which is financially supported by Interreg France-Wallonie-Vlaanderen (Fonds Européen de Développement Régional, FEDER-ERDF). E.V. is financially supported by FRIA (F.R.S.-FNRS). M.L. is financially supported by the WBI Excellence Grant Programme - WBI-World.

## Conflicts of Interest

The authors declare no conflict of interest.

## Author contribution

S.G., M.L. and M.V. conceived the project and S.G. supervised the project. M.L. developed photopolymerization methods of hydrogels. M.V. and M.L performed cell experiments and imaging. M.V. and M.L. and S.G. analyzed data. The article was written by M.L., M.V., E.V. and S.G., read and corrected by all authors, who all contributed to the interpretation of the results.

## Data Availability Statement

The data that support the findings of this study are available in the supplementary material of this article.

---

**Supplementary Figure 1.**
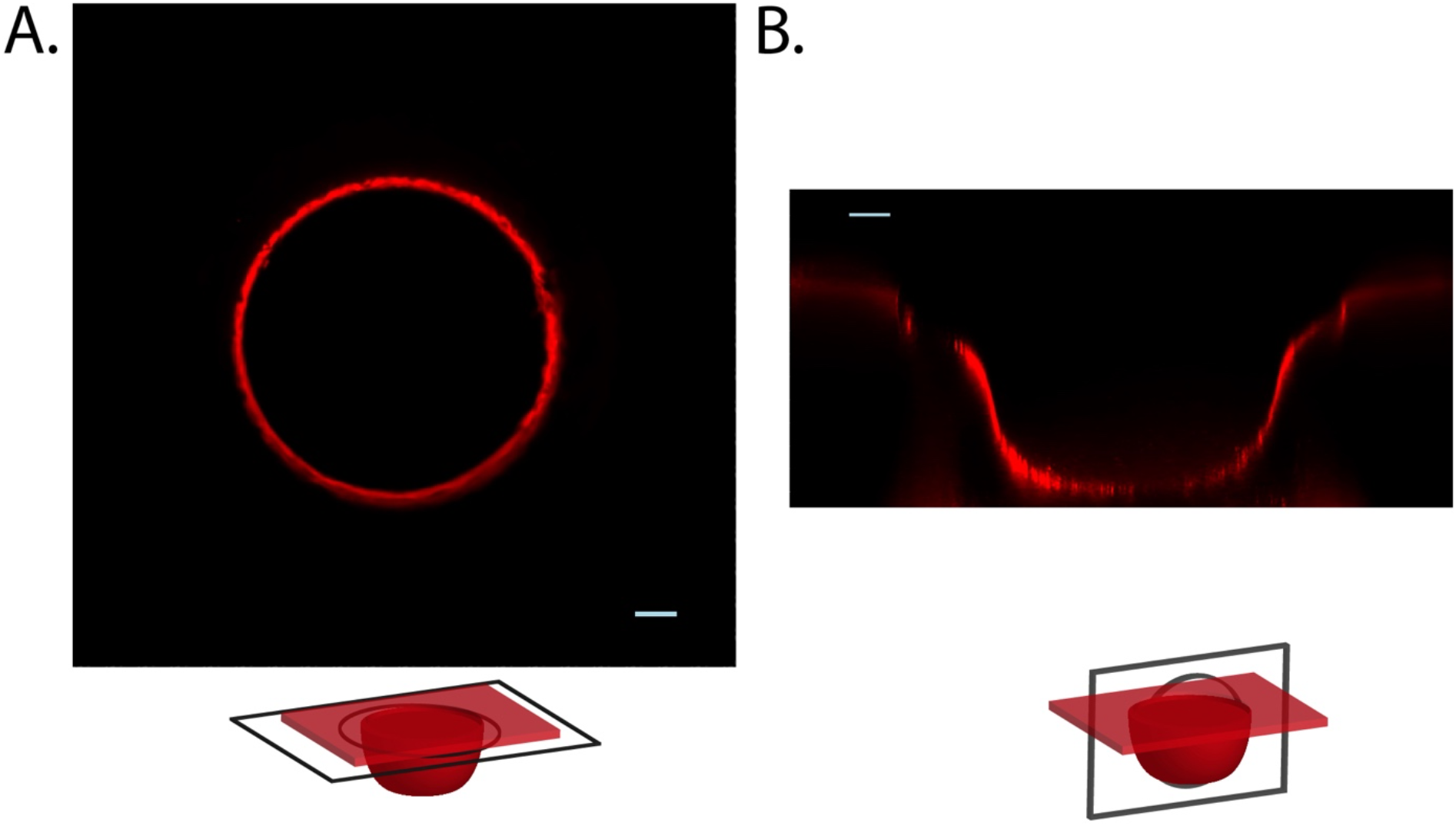
(A) Top and (B) side views of a microwell of 100 μm in radius coated with a uniform layer of human rhodamine fibronectin (in red). The scale bars are 25 μm.

**Supplementary Figure 2.**
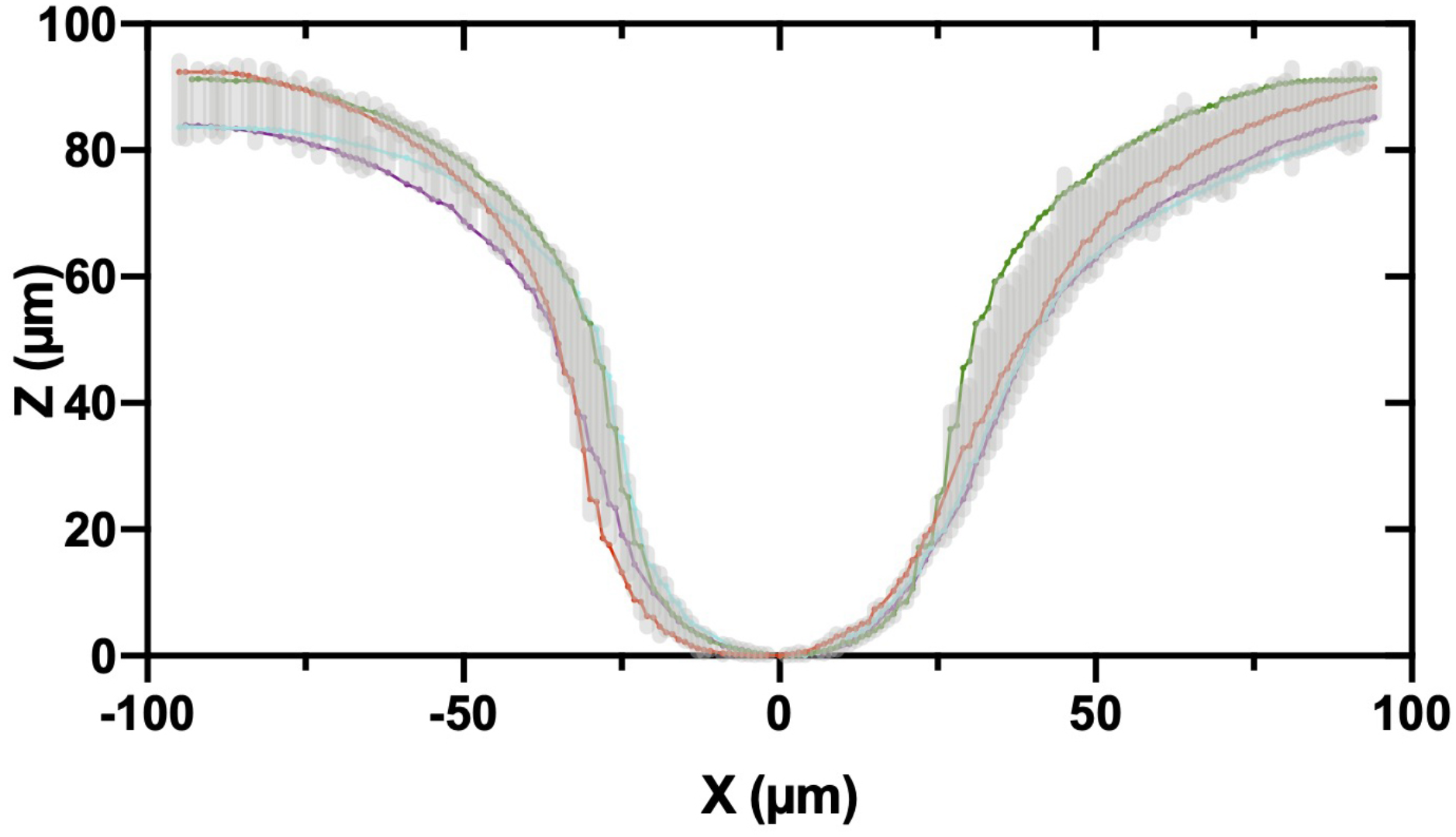
Superimposed (*xz*) profiles of microwells of 65 μm in radius. Each solid curve (n=4 replicates) corresponds to the mean of 3 microwell profiles.

**Supplementary Movie S1 –** 3D confocal view of a microwell of 30 μm in radius covered by an epithelial monolayer stained for F-actin with phalloidin (in green) and DNA with Hoechst 33342 (in blue).

**Supplementary Movie S2 –** 3D confocal view of a microwell of 65 μm in radius covered by an epithelial monolayer stained for F-actin with phalloidin (in green) and DNA with Hoechst 33342 (in blue).

**Supplementary Movie S3 –** 3D confocal view of a microwell of 100 μm in radius covered by an epithelial monolayer stained for F-actin with phalloidin (in green) and DNA with Hoechst 33342 (in blue).

**Supplementary Movie S4 –** Time-lapse movie (duration time = 10 hours) in confocal mode of the nuclear movements in an epithelial monolayer covering a 3D microwell of 100 μm in radius. Nuclei are stained with Hoechst 33342. The scale bar is 50 μm.

## Notes

### Competing Interest Statement

The authors have declared no competing interest.

